# Presynaptically silent synapses are modulated by the density of surrounding astrocytes

**DOI:** 10.1101/2020.03.20.000166

**Authors:** Kohei Oyabu, Kotomi Takeda, Hiroyuki Kawano, Kaori Kubota, Takuya Watanabe, N. Charles Harata, Shutaro Katsurabayashi, Katsunori Iwasaki

## Abstract

The astrocyte, a major glial cell type, is involved in formation and maturation of synapses, and thus contributes to sustainable synaptic transmission between neurons. Given that the animals in the higher phylogenetic tree have brains with higher density of glial cells with respect to neurons, there is a possibility that the relative astrocytic density directly influences synaptic transmission. However, the notion has not been tested thoroughly. Here we addressed it, by using a primary culture preparation where single hippocampal neurons are surrounded by a variable but countable number of cortical astrocytes in dot-patterned microislands, and recording synaptic transmission by patch-clamp electrophysiology. Neurons with a higher astrocytic density showed a higher amplitude of evoked excitatory postsynaptic current (EPSC) than that of neurons with a lower astrocytic density. The size of readily releasable pool of synaptic vesicles per neuron was significantly higher. The frequency of spontaneous synaptic transmission (miniature EPSC) was higher, but the amplitude was unchanged. The number of morphologically identified glutamatergic synapses was unchanged, but the number of functional ones was increased, indicating a lower ratio of presynaptically silent synapses. Taken together, the higher astrocytic density enhanced excitatory synaptic transmission by increasing the number of functional synapses through presynaptic un-silencing.

## INTRODUCTION

While neurons are indispensable for information processing through neural circuits, it has been reported that glial cells play essential roles in brain physiology and development (Barres, 2008). Dynamic information processing in the brain is not only due to the interaction between neurons, but also to the interaction between neurons and astrocytes, a major type of glial cells. In particular, the structure and function of synapses, the basic information-processing units of neurons, are closely regulated by astrocytes. For example, astrocytes are involved in the regulation of synaptic strength (Haydon, 2001) and synapse formation (Nedergaard et al., 2003). Neurons co-cultured with astrocytes have higher synaptic efficacy compared with neurons in the absence of astrocytes, based on direct contact with and humoral factors secreted by the astrocytes (Pfrieger et al., 1997: Ullian et al. 2001: Hama et al., 2004: Crawford et al., 2012: Sobieski, et al, 2015). Furthermore, astrocytes are essential in long-term memory acquisition (Suzuki et al., 2011). These findings suggest that astrocytes perform an essential role in higher brain functions such as memory and learning.

Glial cells and neurons possess one notable evolutionary feature. The ratio of the number of glia to that of neurons in the brain (glia/neuron ratio) is higher for animal species in the higher phylogenetic tree (Nedergaard et al., 2003). For example, the ratio is 0.03-0.05 in the leech ganglia, but it increases to approximately 0.3 in rodent cerebral cortex, and reaches approximately 1.4 in human cerebral cortex. Thus, this evolutionary principle seems to be associated with the complexity of brain functions. The glia/neuron ratio is loosely correlated with the brain size (higher ratios with larger brains), but is tightly and negatively correlated with neuronal density (higher ratios with lower neuronal densities) (Herculano-Houzel, 2009, 2014; von Bartheld et al., 2016). The low neuronal density in human brains is considered to reflect the large size of the neurons (Herculano-Houzel, 2014). In support of these anatomical findings, human cerebral cortical neurons are larger than the rat counterparts, and the longer human dendrites were shown to increase electrical compartmentalization, change the neuronal input-output properties, and therefore influence the synaptic integration and computation (Beaulieu-Laroche, et al, 2018). These findings illustrate that the glia/neuron ratios are linked with key functional differences of neurons, and are important in comparing different animals’ species. However, the glia/neuron ratios also vary in different brain regions of a given species, including humans (Azevedo et al., 2009). The direct impact of such intra-species variations in glia/neuron ratios has been unexplored.

This study thus evaluated the influence of different astrocyte densities on synaptic transmission in a given animal species. This was achieved by co-culturing single mouse neurons on microislands of mouse astrocytes, and recording synaptic transmission in each microisland by patch-clamp electrophysiology. By counting the number of astrocytes, we were able to define the astrocyte/neuron ratio for each microisland. The electrophysiological results were separated into two groups, according to the low or high densities of astrocytes. We show that the excitatory synaptic transmission is enhanced with the increased astrocyte density, through a presynaptic mechanism.

## EXPERIMENTAL PROCEDURES

### Animals

All procedures regarding animal care were performed in strict accordance with the rules of the Experimental Animal Care and Welfare Committee of Fukuoka University, following approval of the experimental protocol (Permit Numbers: 1602907 and 1712128). Timed-pregnant Jcl:ICR mice (Catalogue ID: Jcl:ICR, CLEA Japan, Inc., Tokyo, Japan) were purchased at gestational day 15 from the Kyudo Company (Tosu, Japan). Fifteen to seventeen-week-old pregnant Jcl:ICR mice were used. The bodyweights of the pregnant mice were not recorded. A pregnant mouse was housed individually in a plastic mouse-cage in temperature-controlled rooms (23 ± 2°C) at our animal facility with a 12-hour light-dark cycle. Food (CLEA Rodent Diet, CE-2, CLEA Japan, Inc., Tokyo, Japan) and water were provided *ad libitum*.

### Autaptic neuron culture

Astrocytes and neurons from newborn timed-pregnant Jcl:ICR mice were cultured as described previously (Bekkers and Stevens, 1991; Oyabu et al. 2019). In brief, cerebral cortices were obtained from newborn mice of either sex at postnatal days 0–1. The cerebral cortices were trypsinized and dissociated. Cells were cultured in 75 cm^2^ culture flasks (Corning Inc., Corning, NY, USA). After 2 weeks, non-astrocytic cells were removed by tapping the culture flask several times. After that, astrocytes were isolated and plated at a density of 6,000 cells/ cm^2^ per well onto 22-mm coverslips (thickness no. 1; Matsunami, Osaka, Japan) in 6-well plates (TPP, Switzerland). The coverslips were first coated with 0.5% agarose, and then were stamped with a 1:1 mixture of rattail collagen (final concentration 1.0 mg/mL, BD Biosciences, San Jose, CA, USA) and poly-D-lysine (final concentration 0.25 mg/mL, Sigma-Aldrich, St Louis, MO, USA) in 300-μm square islands. After 1 week, neurons were obtained from the hippocampus of another newborn mouse of either sex at postnatal days 0-1. The dissociated neurons were plated at a density of 1,500 cells/cm^2^ per well onto the micro-island astrocytes.

The cells were cultured in a humidified incubator at 37°C with 5% CO_2_ for 13-18 days before they were used for electrophysiological, immunocytochemical and functional imaging experiments.

### Electrophysiology

Synaptic responses were recorded from autaptic neurons, using a patch-clamp amplifier (Multi-Clamp 700B, Molecular Devices, Sunnyvale, CA, USA), in the whole-cell configuration under the voltage-clamp mode, at a holding potential (Vh) of −70 mV, and room temperature (23 ± 2°C) in all cases. Patch-pipette resistance was 4–5 MΩ, and 70%–90% of access resistance was compensated. Autaptic neurons showed the evoked synaptic transmission in response to an action potential elicited by a brief (2 ms) somatic depolarization pulse (to 0 mV) from the patch pipette. The synaptic responses were recorded at a sampling rate of 20 kHz and were filtered at 10 kHz. Data were excluded from analysis if a leak current of >300 pA was observed. The data were analyzed offline using AxoGraph X 1.2 software (AxoGraph Scientific, Berkeley, CA, USA). Miniature excitatory postsynaptic currents (mEPSCs) were recorded for 100 sec. They were detected with an amplitude threshold of 5 pA, using AxoGraph X 1.2 software. The evoked EPSC and mEPSC were confirmed to be mediated by the excitatory neurotransmitter glutamate, based on an effective block of the EPSCs by CNQX (data not shown)

### Nuclear staining in live cells

Immediately before patch-clamp recording, the nuclei in live cells were stained using NucBlue™ Live ReadyProbes™ reagent (Thermo Fisher Scientific, Waltham, MA, USA). Briefly, two drops of NucBlue Live ReadyProbes Reagent were added to the culture dish per milliliter of medium. The cells were then incubated in a humidified incubator at 37°C with 5% CO_2_ for 20 min, under protection from ambient light. The stained cells were observed on an inverted microscope (Eclipse-Ti2-U, Nikon, Tokyo, Japan), equipped with a 365-nm LED light source (KSL70, Rapp OptoElectronic, Hamburg, Germany) and a filter cube (375/28-nm excitation, 415-nm dichroic long-pass, 460/60-nm emission).

### Immunocytochemistry

Autaptic neurons were immuno-stained, as described previously (Kawano et al., 2012; Oyabu et al. 2019). Primary antibodies were used at the following dilutions: anti-microtubule-associated protein 2 (MAP2), 1:1,000 (guinea pig polyclonal, antiserum, Synaptic Systems, Göttingen, Germany), anti-vesicular glutamate transporter 1 (anti-VGLUT1), 1:2,000 (rabbit polyclonal, affinity-purified, Synaptic Systems). Appropriate secondary antibodies conjugated to Alexa Fluor 488 (anti-guinea pig) or 594 (anti-rabbit) (Thermo Fisher Scientific, Waltham, MA, USA) were used at a dilution of 1:400. Cell nuclei were visualized by counterstaining with DAPI contained in the mounting medium (ProLongH Gold antifade mounting reagent, Thermo Fisher Scientific, Waltham, MA, USA). Autaptic neurons were observed using a confocal microscope (LSM710, Carl Zeiss, Oberkochen, Germany) with a 40× objective lens (C-Apochromat, numerical aperture 1.2) to count the number of excitatory glutamatergic synapses (for Figure 4).

### Identification of presynaptically active synapses using FM1-43

Presynaptic terminals that actively release neurotransmitters, namely, active synapses, were quantified in autaptic neuronal cultures using N-(3-triethylammoniumpropyl)-4-(4-(dibutyl amino) styryl) pyridinium dibromide (FM1-43, Thermo Fisher Scientific, Waltham, MA, USA), similarly as in our previous report (Kawano et al., 2012). Briefly, the synaptic recycling vesicles were loaded with 10 μM FM1-43 in a high-potassium (45 mM) extracellular solution containing the NMDA receptor antagonist (2R)-amino-5-phosphonovaleric acid (APV, 25 μM, Sigma-Aldrich, St Louis, MO, USA) and the AMPA receptor antagonist 6-cyano-7-nitroquinoxaline-2,3-dione (CNQX, 10 μM, Sigma Aldrich), for 2 min at room temperature. The coverslip was washed three times for 2 minutes each with a standard extracellular solution containing 1 μM tetrodotoxin (TTX), a Na^+^ channel blocker, to remove excess FM1-43. Autaptic neurons were then fixed in a 4% paraformaldehyde solution of phosphate-buffered saline (PBS) for 10 min. To minimize the loss FM1-43 signals (e.g. by photobleaching by ambient light), the images were captured soon after the neurons were fixed. Sixteen-bit images were acquired with a scientific CMOS camera (pco.edge 4.2, pco, Kelheim, Germany) on an inverted microscope (Eclipse-TiE, Nikon, Tokyo, Japan) with a 40× objective lens (Plan Apoλ, numerical aperture 0.95). FM1-43 was excited using a white LED (Lambda HPX, Sutter Instruments, Novato, CA, USA) at 100% of maximum intensity, and imaged using a filter cube (470/40-nm excitation, 595-nm dichroic long-pass, 535/50-nm emission). In each sample, ten images were captured with the exposure time of 300 ms per image, averaged, and used for analysis based on the average-intensity of the pixels.

### Identification of presynaptically silent synapses

After taking images of FM1-43 puncta, the fixed autaptic neuron was blocked and permeabilized with PBS containing 5% normal goat serum and 0.1% Triton X-100 for 30 min in the microscope chamber. After blocking and permeabilizing, it was confirmed that the FM1-43 puncta were completely de-stained (data not shown). The samples were then immuno-stained for VGLUT1 and MAP2 as described above. This step was also performed in the microscope chamber (Kawano et al., in preparation). Although the emission spectrum of Alexa Fluor 488 considerably overlapped that of FM1-43, the acquisition of one signal did not interfere with the acquisition of the other. This was because the FM1-43 signal was lost completely by treatment with 0.1% Triton X-100, which was necessary for immuno-staining procedure before Alexa Fluor 488 signal was acquired. Alexa Fluor 488 (for MAP2) was therefore imaged using the same optical system as for the FM1-43. Alexa Fluor 594 (for VGLUT1) was excited using a white LED at 100% of maximum intensity, and imaged using a filter cube (560/40-nm excitation, 595-nm dichroic long-pass, 630/60-nm emission). Ten images per sample were captured as for the FM1-43 puncta, with the exposure time of 300 ms per image, and then averaged.

To identify presynaptically silent synapses, the image of FM1-43 was merged with that of VGLUT1 and MAP2 using ImageJ (1.48v, Wayne Rasband, NIH, available at http://imagej.nih.gov/ij/). For this purpose, the grayscale images of MAP2, FM1-43 and VGLUT1were converted to pseudo-color images in blue, green and red, respectively (for Figure 5). The VGLUT1 puncta that were not stained with FM1-43 were defined as presynaptically silent synapses.

### Solutions

The standard extracellular solution was: 140 mM NaCl, 2.4 mM KCl, 2 mM CaCl_2_, 1 mM MgCl_2_, 10 mM glucose, 10 mM HEPES (pH 7.4, 320 mOsm). The extracellular solution for application of FM1-43 was: 97.4 mM NaCl, 45 mM KCl, 2 mM CaCl_2_, 1 mM MgCl_2_, 10 mM glucose, 10 mM HEPES (pH 7.3, 320 mOsm). Patch pipettes were filled with an intracellular solution (146.3 mM K-gluconate, 0.6 mM MgCl_2_, 2.4 mM ATP-Na_2_, 0.3 mM GTP-Na_2_, 50 U/ml creatine phosphokinase, 12 mM phosphocreatine, 1 mM EGTA, 17.8 mM HEPES, pH 7.4). Miniature excitatory postsynaptic currents (mEPSCs) were recorded using the standard extracellular solution containing 1 μM TTX. Hypertonic solutions for determining the size of the readily releasable pool from synaptic vesicles (RRP) were prepared by adding 0.5 M sucrose to the standard extracellular solution. The extracellular solutions were applied using a fast-flow application system (SF-77B, Warner Instruments, Hamden, CT, USA). Each flow pipe has a large diameter (430 μm), ensuring that the solution is applied to all parts of an autaptic neuron on an astrocytic micro island (300 × 300 μm). This configuration was suitable for the application of sucrose to induce synaptic responses from all nerve terminals of the recorded neuron. All chemicals were purchased from Sigma Aldrich (St Louis, MO, USA), except where otherwise specified.

### Statistical analysis

Data were expressed as the mean ± SEM. Statistical analysis was performed using Student’s unpaired t-test for the comparison of two groups. Significance was considered when *p <* 0.05.

## RESULTS

### Excitatory synaptic transmission is enhanced with high density of astrocytes

We used autaptic cultures (Bekkers et al. 1991, Oyabu et al. 2019) to assess whether a difference in astrocyte densities around neurons affects synaptic transmission. This preparation has two major advantages. First, it allows single neurons to be plated on variable numbers of astrocytes per microisland. By combining this advantage with live-cell nuclear staining, we can count the number of astrocytes in each microisland and classify the synaptic transmission phenotypes based on the astrocytic density. Second, the neurotransmitter release from any nerve terminal of a single identified neuron can be detected by patch-clamp electrophysiological recording, albeit with different sensitivities due to different distances of postsynaptic receptors from the recording electrode. Figure 1 shows representative images of phase-contrast optics and nuclear staining of microislands with low (Fig. 1A-C) and high numbers of astrocytes (Fig. 1D-F). Because we used the microislands that contained only single neurons, the ratio of the number of astrocytes to that of neurons (astrocyte/neuron) is simply the number of astrocytes in each dot-patterned microisland. In this study, we classified the microislands into two broad experimental groups: those with astrocyte/neuron = 1 to 10 were defined as a low-density group (LDG, Fig. 1A-C), and those with astrocyte/neuron = 20 to 30 were defined as a high-density group (HDG, Fig. 1D-F). All the subsequent analyses were based on the comparison between these two groups which were cultured in the same culture dishes.

**Figure 1.**
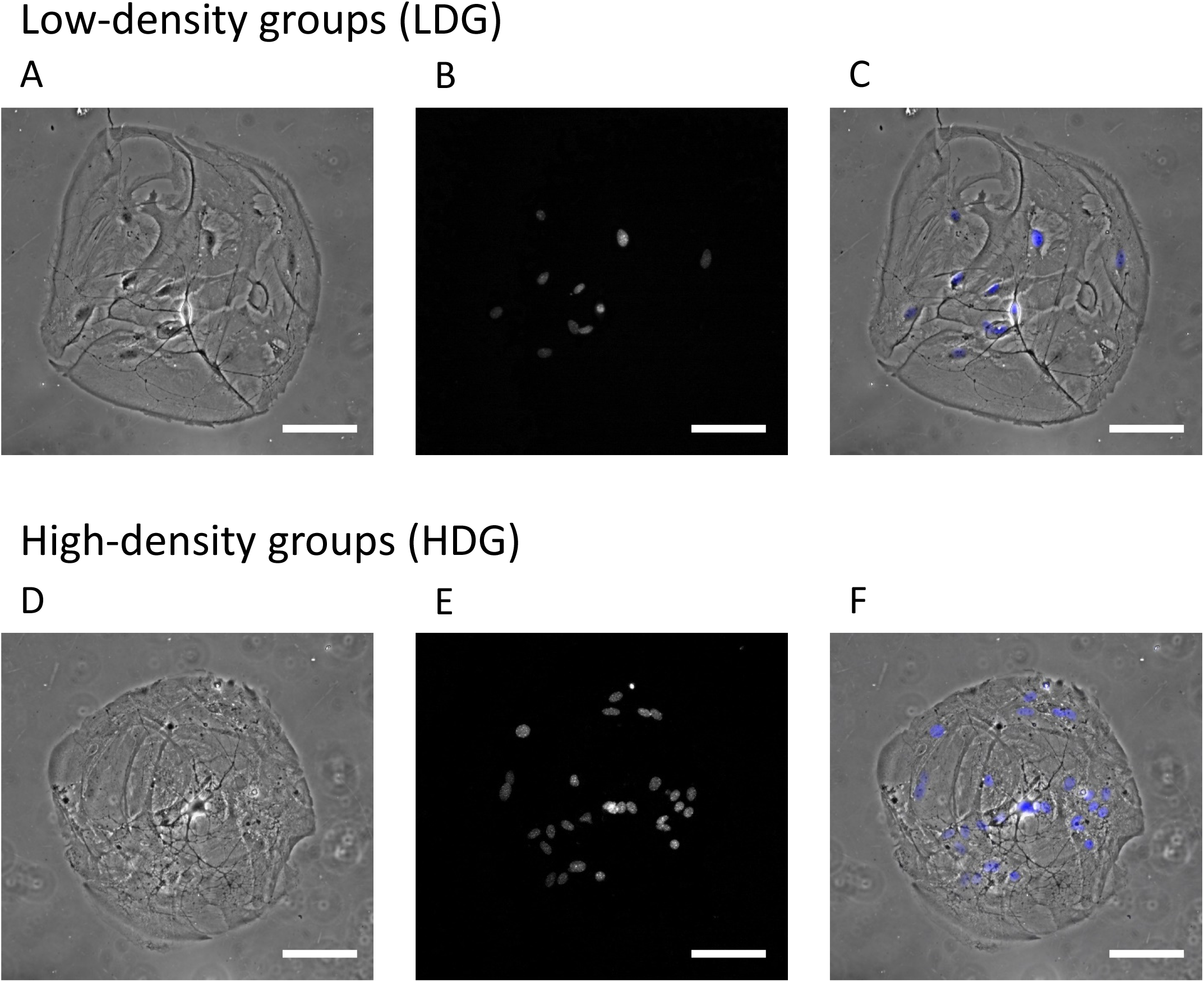
Single autaptic hippocampal neurons co-cultured with different numbers of astrocytes. (A) A representative phase-contrast image of an autaptic neuron cultured with a low density of astrocytes. (B) Nuclear staining of live astrocytes and a single neuron shown in (A). (C) A merged image of phase-contrast and nuclear staining images of the same observation field. (D) A representative phase-contrast image of an autaptic neuron cultured with a high density of astrocytes. (E) Nuclear staining of live astrocytes and a single neuron shown in (D). (F) A merged image of phase-contrast and nuclear staining images of the same observation field. All scale bars indicate 100 μm.

We recorded and compared the evoked excitatory postsynaptic currents (EPSCs) in LDG and HDG (Fig. 2A). The evoked EPSC amplitude was significantly larger in the HDG than that of the LDG (LDG, 4.62 ± 0.53 nA; HDG, 7.12 ± 0.71 nA; Fig. 2B). Next, mEPSCs were recorded in the presence of TTX (Fig. 2C). Since mEPSCs correspond to the activation of postsynaptic receptors by neurotransmitters spontaneously released from single synaptic vesicles (Katz, 1979. Bekkers, 1995), it is commonly understood that the changes in mEPSC frequencies mostly reflect the changes in the number of functional nerve terminals or the probability of release from nerve terminals, whereas the changes in mEPSC amplitudes mostly reflect those in the number or properties of postsynaptic receptors. We found that the mEPSC frequency was significantly higher in the HDG than that of the LDG (LDG, 7.07 ± 0.84 Hz; HDG, 9.88 ± 1.01 Hz; Fig. 2D), whereas the mEPSC amplitude was identical in both groups (LDG, 29.1 ± 1.01 pA; HDG, 29.0 ± 1.43 pA; Fig. 2E). These results show increases in the evoked EPSC amplitude and mEPSC frequency by high-density astrocytes, and imply that these increases in excitatory synaptic transmission originated from changes in presynaptic properties.

**Figure 2.**
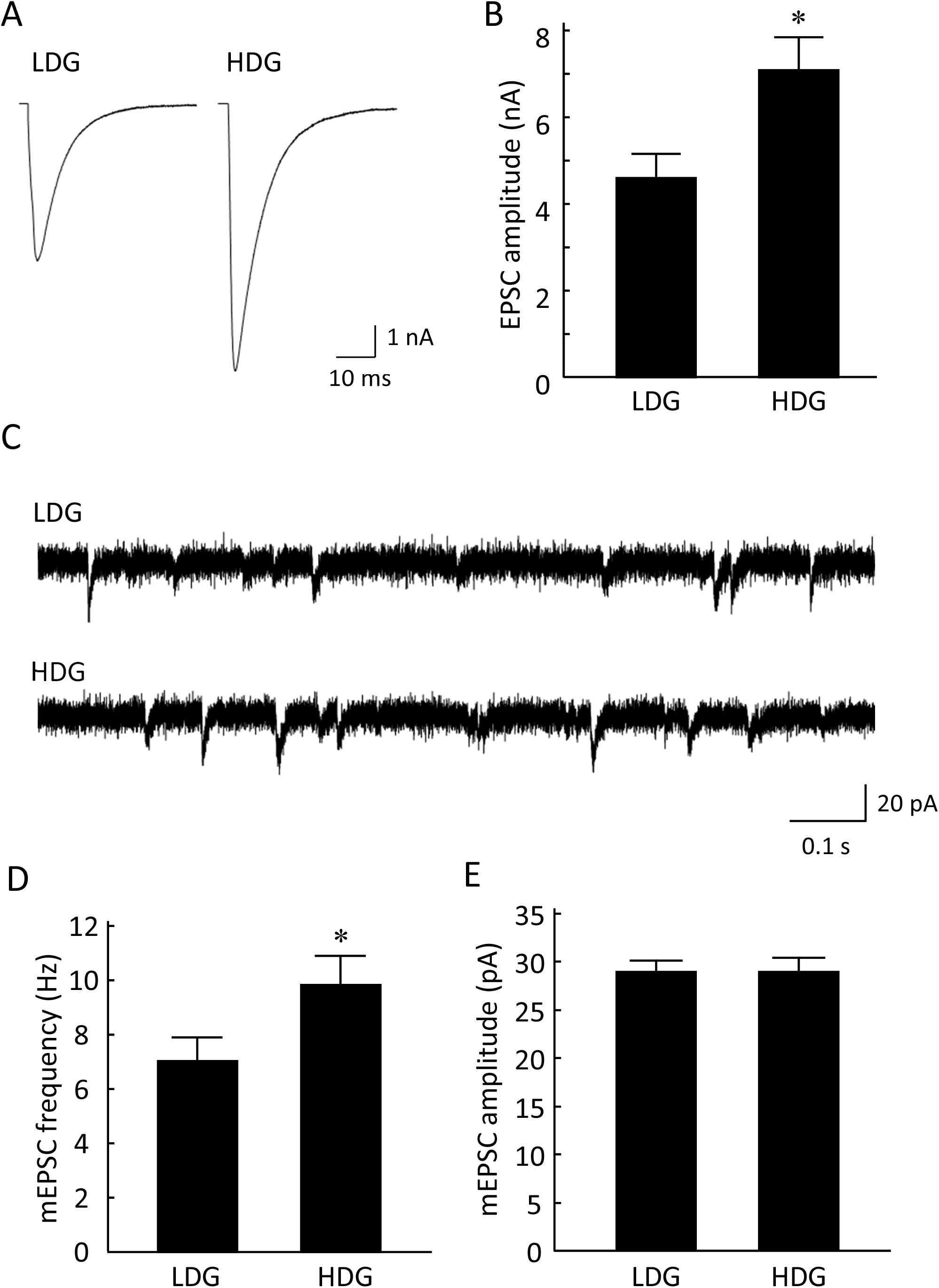
Excitatory synaptic transmission was enhanced with the increased astrocyte density. (A) Representative traces of evoked excitatory postsynaptic currents (EPSCs) recorded electrophysiologically from autaptic neurons (LDG and HDG). Averaged trace of eight stimuli at 0.2 Hz is shown. Depolarization-induced action currents have been removed for clarity. (B) Average amplitudes of evoked EPSCs in autaptic neurons co‐cultured with LDG (n = 66 neurons /10 cultures) or HDG (n = 60 neurons /10 cultures). (C) Representative traces of miniature EPSCs (mEPSCs) in the LDG or HDG. (D) The frequency of mEPSCs in the LDG (n = 66 neurons /10 cultures) or HDG (n = 60 neurons /10 cultures). (E) The amplitude of mEPSCs in the LDG (n = 66 neurons /10 cultures) or HDG (n = 60 neurons /10 cultures). *p < 0.05.

### Astrocyte density differences do not change synaptic release machineries and synapse formation

Because our data showed that the high astrocyte density affected presynaptic properties, we measured the size of the readily releasable pool (RRP) of synaptic vesicles (Fig. 3A). The RRP size was determined by the application of hypertonic 0.5 M sucrose solution to a single microisland (Rosenmund and Stevens, 1996; Kawano et al, 2012). RRP size was significantly increased in HDG (LDG, 0.96 ± 0.15 nC; HDG, 1.38 ± 0.15 nC; Fig. 3B). Since all the functional nerve terminals in a microisland contribute to the RRP measurement by our method, the increase in RRP may be due to an increase in the functions of individual nerve terminals or the number of nerve terminals in each autaptic neuron.

**Figure 3.**
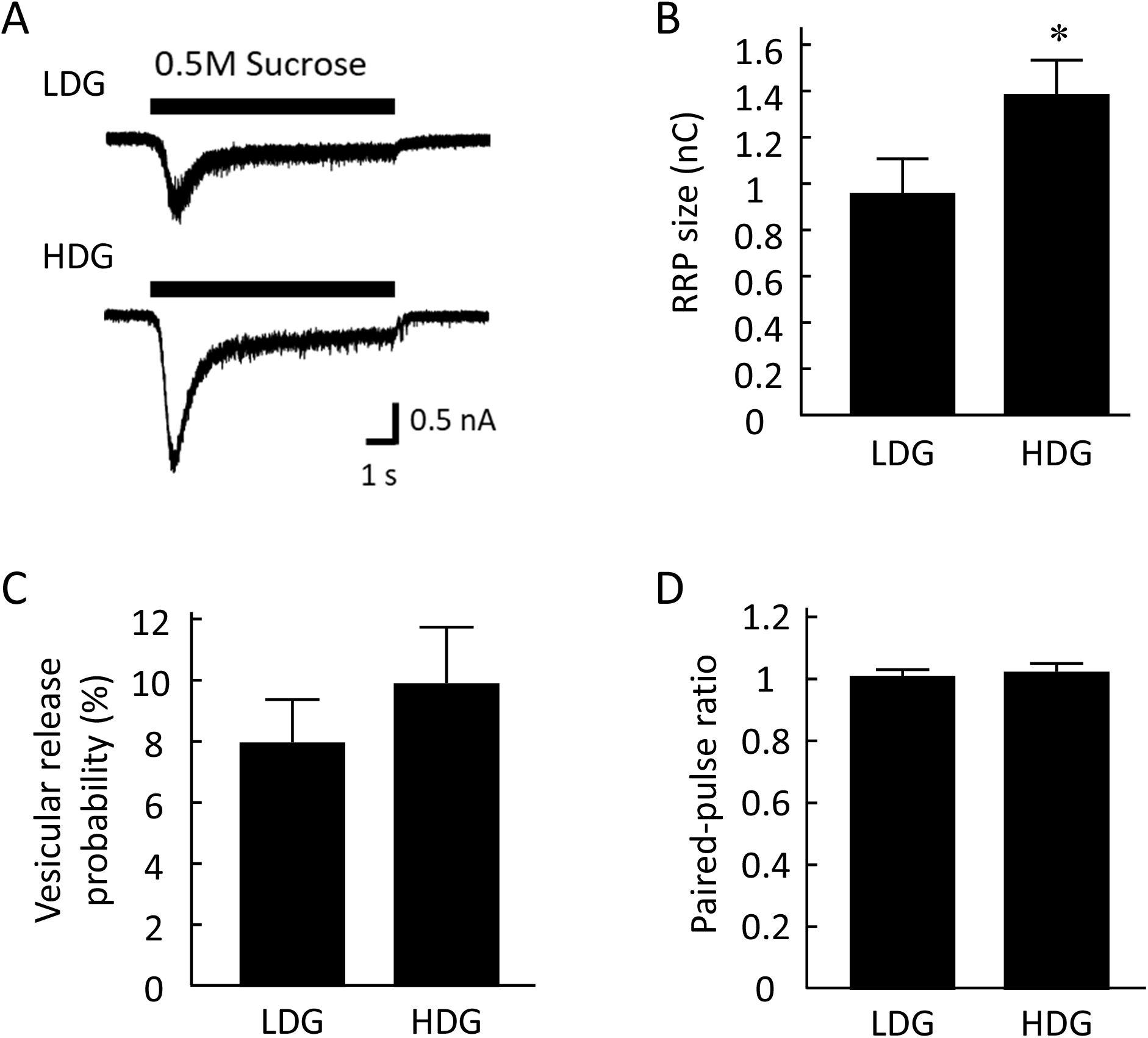
Synaptic release machinery was unaltered by differences in astrocyte density. (A) Representative traces of the responses to 0.5 M sucrose solution (8 sec) in the LDG or HDG. (B) The averaged size of the readily releasable pool (RRP), measured by the response to sucrose, in the LDG (n = 66 neurons /10 cultures) or HDG (n = 60 neurons /10 cultures). (C) The vesicular release probability in the LDG (n = 66 neurons /10 cultures) or HDG (n = 60 neurons /10 cultures). (D) The paired‐pulse ratio (EPSC_2_ / EPSC_1_) in the LDG (n = 46 neurons /7 cultures) or HDG (n = 42 neurons /7 cultures). *p < 0.05.

In order to further examine presynaptic functions, we designed two experiments. First, we analyzed the vesicular release probability (Pvr). Pvr was defined as the fraction of vesicles releasable by an action potential among the RRP vesicles. It was calculated by dividing the electric charge of an action potential–induced EPSC (i.e. area of the evoked EPSC trace) by the electric charge of sucrose-induced transient EPSC (i.e. area of the transient component of the trace). Pvr was not statistically different between the two groups (LDG, 7.98 ± 1.39%; HDG, 9.91 ± 1.83%; Fig. 3C). Second, we measured the paired-pulse ratio of evoked EPSCs, by giving two depolarizing pulses (action potentials) with an inter-pulse interval of 50 ms. This parameter was calculated as the amplitude ratio of the second to the first response (EPSC_2_ / EPSC_1_). This parameter is inversely correlated with the probability of release, and is generally thought to reflect multiple factors, such as the amount of Ca^2+^ influx following an action potential, kinetics of the increase in cytosolic Ca^2+^ concentration, and the availability or depletion of synaptic vesicles for release (Xu-Friedman and Regehr, 2004). The paired-pulse ratio was not different between the two groups (LDG, 1.00 ± 0.02; HDG, 1.02 ± 0.03; Fig. 3D). A lack of changes in the Pvr or the paired-pulse ratio indicates that the astrocyte density did not affect the presynaptic functions, specifically the properties associated with neurotransmitter release from functional or active nerve terminals.

Because functions of individual nerve terminals did not change, the increased excitatory synaptic transmission may have resulted from an increased number of synapses. Nerve terminals of excitatory neurons in the hippocampus predominantly express the vesicular glutamate transporter 1 (VGLUT1) (Wojcik et al., 2004). Therefore, we identified the excitatory nerve terminals as the punctate structures stained positively with VGLUT1, and counted their number. Surprisingly, the numbers were not different between the two groups (LDG, 358.29 ± 32.32; HDG, 412.21 ± 43.29; Fig. 4B). Thus, in the HDG group, there was enhancement of functions of the collective nerve terminals (Figs. 2D, 3B), whereas there was no change in the functions of the individual nerve terminals that released neurotransmitters (Figs. 3C, D) or in the numbers of morphologically identified terminals (Fig. 4).

**Figure 4.**
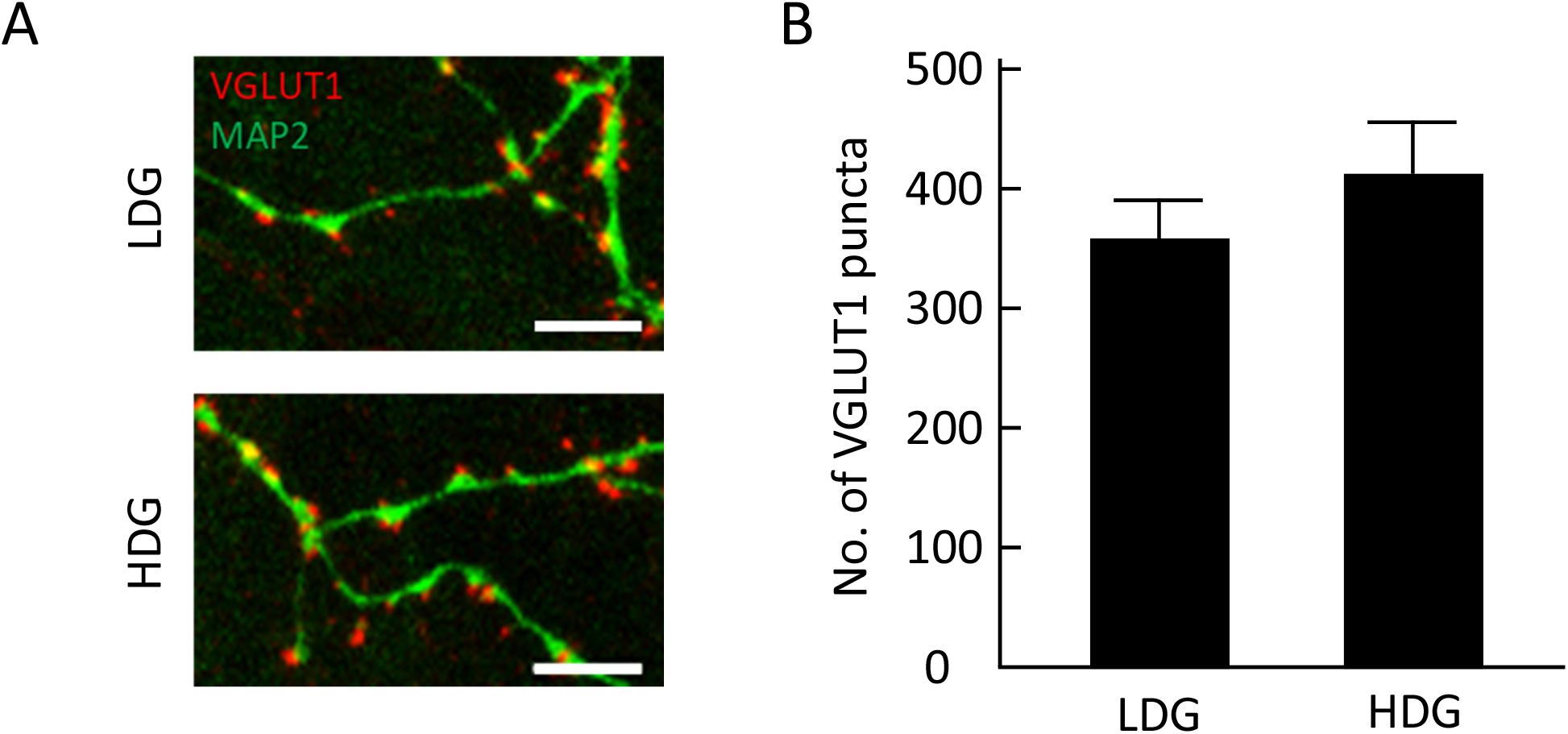
Differences in astrocyte density did not affect the number of excitatory synapses. (A) Representative images of autaptic neurons immuno-stained for the dendritic marker, microtubule‐associated protein 2 (MAP2) (in green) and the excitatory synapse marker, vesicular glutamate transporter 1 (VGLUT1) (in red). The scale bars indicate 10 μm. (B) The number of VGLUT1 puncta in the LDG (n = 31 neurons /4 cultures) or HDG (n = 24 neurons /4 cultures).

### The ratio of presynaptically silent synapses was reduced by increasing the astrocyte density

The above results are consistent with the concept that the astrocytic density modified presynaptically silent synapses. Presynaptically silent synapses are morphologically mature, but their synaptic vesicles do not exocytose (i.e. they do not release neurotransmitters) at all even when challenged with a robust depolarizing stimulus or Ca^2+^influx (Crawford and Mennerick, 2012). In other words, the presynaptically silent synapses are not detectable electrophysiologically, and therefore they do not contribute to the release parameters, such as EPSCs, RRP and Pvr. We therefore posited that the synaptic transmission was enhanced by a decreased ratio of presynaptically silent synapses in the HDG.

To test this hypothesis directly, we identified the presynaptically silent synapses, by using double staining with FM1-43 and VGLUT1. In this experiment, FM1-43 was used to locate functional nerve terminals by loading into recycling synaptic vesicles that undergo endocytosis after stimulus-induced exocytosis, while the VGLUT1 immunostaining was used to locate mature nerve terminals morphologically regardless of the capability of endocytosis. In previous studies, synapses labeled with VGLUT1 but lacking FM1–43 loading were identified as presynaptically silent synapses (Moulder et al., 2004; Kawano et al., 2012). The ratio of presynaptically silent synapses (VGLUT1 +, FM1–43 −) among all excitatory synapses (VGLUT1 +) was significantly smaller in the HDG (LDG, 26.81 ± 3.68%; HDG, 20.35 ± 1.86%; Fig. 5B). This result supports the hypothesis that the astrocytic density influences presynaptically silent synapses, such that the number of functional nerve terminals is increased by high astrocytic density.

**Figure 5.**
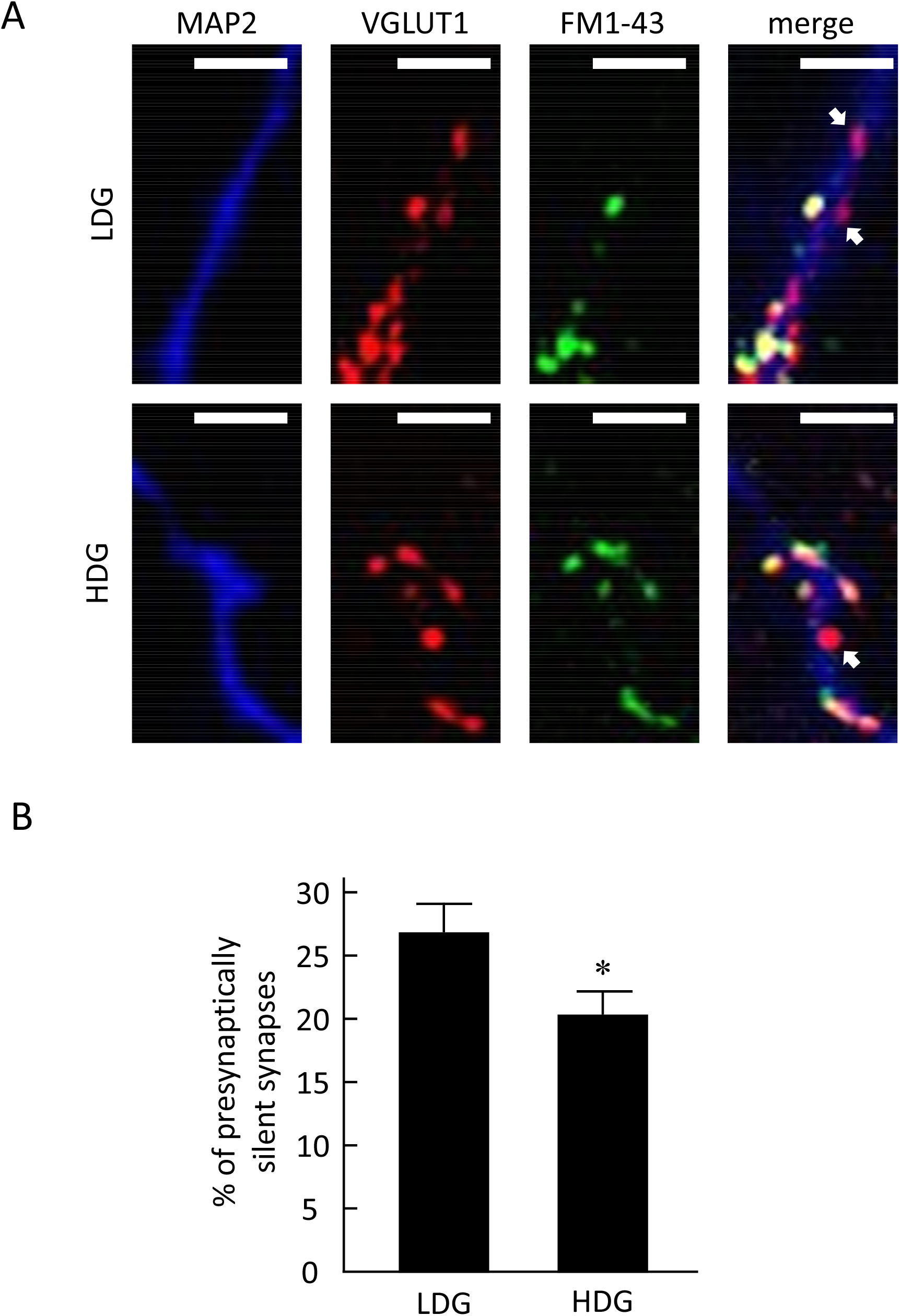
Ratio of presynaptically silent synapses is reduced by increasing astrocyte density. (A) Images of MAP2 immunostaining (in blue), VGLUT1 immunostaining (in red), and FM1-43 labeling of presynaptic terminals (in green), in LDG (top) and HDG (bottom). In the merged images, the arrows indicate presynaptically silent synapses, stained positively for VGLUT1 but negatively for FM1-43. All scale bars indicate 5 μm. (B) The ratio of presynaptically silent synapses in the LDG (n = 13 neurons / 11cultures) or HDG (n = 13 neurons /11 cultures), *p < 0.05.

## DISCUSSION

In this study, we evaluated the impact of astrocytic density on excitatory synaptic transmission, using single autaptic neurons that were co-cultured with astrocytes at different densities. The HDG exhibited significant increases in the factors that reflect the collective properties of functional nerve terminals in our experimental system: the evoked EPSC amplitude (Fig. 2B), mEPSC frequency (Fig. 2D), and RRP size (Fig. 3B). In contrast, there was no change in the functions of individual nerve terminals that released neurotransmitters (i.e. functions of presynaptically active synapses): the vesicle release probability (Fig. 3C) and the paired-pulse ratio (Fig. 3D). There was no change in a postsynaptic factor: the mEPSC amplitude (Fig. 2E). Considering an unchanged number of morphologically identified, excitatory glutamatergic nerve terminals (Fig. 4B), all these features introduced by different astrocyte densities can be explained by a change in a single factor: the ratio of presynaptically silent synapses (Fig. 5B). In the HDG, this factor was significantly decreased, i.e. the number of functional nerve terminals was increased due to un-silencing or awakening of silent ones (Crawford and Mennerick, 2012). This change led to an enhancement of the evoked and miniature excitatory synaptic transmission (Fig. 2B, C), without a change in the functions of individual nerve terminals. Increase in the collective properties can be interpreted to be due to the increased number of functional nerve terminals, each of which did not change any release property. Naturally, there can be more complex scenarios, e.g. the functional properties of unsilenced and the already active nerve terminals changed in opposite directions and canceled each other, leading to the apparent lack of functional changes. However, the original scenario shown above is the simplest one and is compatible with every finding in our system.

Overall, this study underscores the important roles of astrocyte/neuron ratio in regulating presynaptic features of synaptic transmission. When synaptic transmission had been recorded from neurons co-cultured with astrocytes, little attention had been paid to the density of astrocytes surrounding neurons. This study prompts us to exercise caution in interpreting the efficiency of synaptic transmission under multiple experimental conditions. In combination with the experiments where astrocytes were deficient (Pfrieger et al., 1997: Ullian et al. 2001: Hama et al., 2004: Crawford et al., 2012: Sobieski, et al, 2015), our study indicates that astrocytic densities have a wide spectrum of effects on synaptic transmission.

Changes in the neuronal activity have been reported to affect the ratio of presynaptically silent synapses, as a part of homeostatic synaptic plasticity to counteract the initial changes. For example, prolonged depolarization of cultured hippocampal neurons increases presynaptically silent synapses (Moulder et al., 2004). This change was accompanied by a reduction in the evoked EPSC amplitude, mEPSC frequency, and RRP size (in a manner opposite in polarity to our findings), and by no change in the mEPSC amplitude or the number of morphologically identified glutamatergic nerve terminals (similar to our findings). Conversely, a blockade of action potentials with TTX reduces the presynaptically silent synapses in the same preparation (Moulder et al., 2006). The cAMP signaling cascade is involved in both presynaptic terminal silencing and un-silencing. For example, increasing cAMP signaling by forskolin reduces the ratio of presynaptically silent synapses at rest, whereas decreasing it by Rp-cAMPS increases the ratio (Moulder et al., 2008).

It is unclear how the presynaptically silent synapses are modified by different astrocytic densities. One pioneering work in this aspect evaluated the effect of astrocyte-derived soluble factors in inducing the presynaptically silent synapses (Crawford et al., 2012). The authors compared the autaptic synaptic transmission when the neurons were plated on “astrocyte-rich” standard microislands or “astrocyte-poor or -deficient” microislands prepared by chemical fixation of only astrocytes. They identified thrombospondins as the potential secreted agent that induces presynaptic silencing through the cAMP signaling cascade. However, secreted agents are not expected to be responsible for the results in our system, because the two density groups were cultured together in the same culture dish, and thus they must have shared the astrocyte-derived secreted agents. At least one possibility can be speculated, still based on the cAMP-dependent mechanisms described above. The astrocyte-neuron lactate shuttle (ANLS) proposed by Pellerin and Magistretti (1994) may be involved in the reduction of presynaptically silent synapses in our study. Lactic acid produced by astrocyte glycolysis is supplied to neurons where it is converted to pyruvate, which is then metabolized in the mitochondria via oxidative phosphorylation to produce ATP. In the HDG of our study, activation of ANLS may increase the energy supply to neurons, activate the cAMP signaling pathway, and may reduce the ratio of presynaptically silent synapses. Clearly, mechanistic insights into the effect of different astrocytic densities await further studies.

The roles that the presynaptically silent synapses play in regulating the function of the physiological and pathological neuronal network still remain under intense investigations (Crawford and Mennerick, 2012). We propose that the astrocyte density also contributes to the regulation of the ratio of presynaptically silent synapses. In this context, it is of interest to note that the glia/neuron ratio is approximately 1.5 in the gray matter of human prefrontal cerebral cortex (von Bartheld et al., 2016), but the ratio varies drastically across brain regions. For example, the glia/neuron ratio is approximately 0.2 in the cerebellum, 3.8 in the cerebral cortex, and 11.4 in the rest of the human brain (Azevedo et al., 2009). Thus, the astrocyte/neuron ratio can be involved in brain-regional difference of information processing through synaptic transmission. Clarifying how the astrocytes are involved in regulating presynaptically silent synapses could be essential for understanding the functions of the neuronal network and the brain.

## Abbreviations

ANLS: astrocyte-neuron lactate shuttle
APV: (2R)-amino-5-phosphonovaleric acid
CNQX: 6-cyano-7-nitroquinoxaline-2,3-dione
DAPI: 4’,6-diamidino-2-phenylindole
EPSC: excitatory postsynaptic current
FM1-43: N-(3-triethylammoniumpropyl)-4-(4-(dibutyl amino) styryl) pyridinium dibromide
HDG: high-density group
LDG: low-density group
LED: light-emitting diode
MAP2: microtubule-associated protein 2
mEPSC: miniature excitatory postsynaptic current
PBS: phosphate-buffered saline
Pvr: vesicular release probability
RRP: readily releasable pool
TTX: tetrodotoxin
VGLUT1: vesicular glutamate transporter 1
Vh: holding potential

## ACKNOWLEDGMENTS

This work was supported by The United States Department of Defense to N.C.H. (W81XWH-14-1-0301), and by a KAKENHI Grant-in-Aid for Scientific Research (C) to S.K. (No. 17K08328) from the Japan Society for the Promotion of Science. We thank Edanz Group (www.edanzediting.com/ac) for editing a draft of this manuscript.

## REFERENCES

Barres BA. The mystery and magic of glia: A perspective on their roles in health and disease. Neuron. 2008;60:430–440.

Haydon PG. GLIA: listening and talking to the synapse. Nat Rev Neurosci. 2001;2:185–193.

Nedergaard M, Ransom B, Goldman SA. New roles for astrocytes: Redefining the functional architecture of the brain. Trends Neurosci. 2003;26:523–530.

Pfrieger FW, Barres BA. Synaptic efficacy enhanced by glial cells in vitro. Science. 1997;277:1684–1687.

Ullian EM, Sapperstein SK, Christopherson KS, et al. Control of synapse number by glia. Science. 2001;291:657–661.

Hama H, Hara C, Yamaguchi K, et al. PKC signaling mediates global enhancement of excitatory synaptogenesis in neurons triggered by local contact with astrocytes. Neuron. 2004;41:405–415.

Crawford DC, Jiang X, Taylor A, et al. Astrocyte-derived thrombospondins mediate the development of hippocampal presynaptic plasticity in vitro. J Neurosci. 2012;32:13100–13110.

Sobieski C, Jiang X, Crawford DC, et al. Loss of Local Astrocyte Support Disrupts Action Potential Propagation and Glutamate Release Synchrony from Unmyelinated Hippocampal Axon Terminals In Vitro. J Neurosci. 2015; 35:11105–11117.

Suzuki A, Stern SA, Bozdagi O, et al. Astrocyte‐neuron lactate transport is required for long‐term memory formation. Cell. 2011;144:810–823.

Herculano-Houzel S. The human brain in numbers: a linearly scaled-up primate brain. Front Hum Neurosci. 2009;3:31.

Herculano-Houzel S. The glia/neuron ratio: how it varies uniformly across brain structures and species and what that means for brain physiology and evolution. Glia. 2014;62:1377–1391.

von Bartheld CS, Bahney J, Herculano-Houzel S. The search for true numbers of neurons and glial cells in the human brain: A review of 150 years of cell counting. J Comp Neurol. 2016;524:3865–3895.

Beaulieu-Laroche L, Toloza EHS, van der Goes MS et al., Enhanced Dendritic Compartmentalization in Human Cortical Neurons. Cell. 2018;175:643–651.

Azevedo FA, Carvalho LR, Grinberg LT, et al. Equal numbers of neuronal and nonneuronal cells make the human brain an isometrically scaled-up primate brain. J Comp Neurol. 2009;513:532–541.

Bekkers JM, Stevens CF. Excitatory and inhibitory autaptic currents in isolated hippocampal neurons maintained in cell culture. Proc Natl Acad Sci USA. 1991;88:7834–7838.

Oyabu K, Kiyota H, Kubota K, et al. Hippocampal neurons in direct contact with astrocytes exposed to amyloid β25-35 exhibit reduced excitatory synaptic transmission. IBRO Rep. 2019;7:34–41.

Kawano H, Katsurabayashi S, Kakazu Y, et al. Long-term culture of astrocytes attenuates the readily releasable pool of synaptic vesicles. PLoS One. 2012;7:e48034.

Bekkers JM, Stevens CF. Quantal analysis of EPSCs recorded from small numbers of synapses in hippocampal cultures. J Neurophysiol. 1995;73:1145–1156.

Katz B, Miledi R. Estimates of quantal content during ‘chemical potentiation’ of transmitter release. Proc R Soc Lond B Biol Sci. 1979;205:369–378.

Rosenmund C, Stevens CF. Definition of the readily releasable pool of vesicles at hippocampal synapses. Neuron. 1996;16:1197–1207.

Xu-Friedman MA, Regehr WG. Structural contributions to short-term synaptic plasticity. Physiol Rev. 2004;84:69–85.

Wojcik SM, Rhee JS, Herzog E, et al. An essential role for vesicular glutamate transporter 1 (VGLUT1) in postnatal development and control of quantal size. Proc Natl Acad Sci USA. 2004;101:7158–7163.

Crawford DC, Mennerick S. Presynaptically silent synapses: dormancy and awakening of presynaptic vesicle release. Neuroscientist. 2012;18:216–223.

Moulder KL, Meeks JP, Shute AA, et al. Plastic elimination of functional glutamate release sites by depolarization. Neuron. 2004;42:423–435.

Moulder KL, Jiang X, Taylor AA, et al. Physiological activity depresses synaptic function through an effect on vesicle priming. J Neurosci. 2006;26:6618–6626.

Moulder KL, Jiang X, Chang C, et al. A specific role for Ca^2+^-dependent adenylyl cyclases in recovery from adaptive presynaptic silencing. J Neurosci. 2008;28:5159–5168.

Pellerin L, Magistretti PJ. Glutamate uptake into astrocytes stimulates aerobic glycolysis: a mechanism coupling neuronal activity to glucose utilization. Proc Natl Acad Sci USA. 1994;91:10625–10629.

